# PD-1 blockade enhances HIV-1 vaccine-induced CD8⁺ T-cell responses in PWH early ART-treated

**DOI:** 10.1101/2024.07.05.601669

**Authors:** Miguel Marin, Alba Ruiz, Esther Jimenez-Moyano, Dan Ouchi, Oscar Blanch-Lombarte, Ruth Peña, Dan Gorman, Richard Barnard, Tomas Hanke, Bonaventura Clotet, Bonnie Howell, Christian Brander, Beatriz Mothe, Julia G Prado

## Abstract

Current HIV-1 therapeutic vaccines do not generate T-cell immune responses in people with HIV-1 (PWH) able to eradicate the virus or to broadly control viral replication. The expression of inhibitory receptors (IRs) may limit vaccine efficacy by reducing specific effector functions in vaccine-induced T-cells. Recently, the clinical use of antibodies targeting IRs or immune checkpoint blockade (ICB) has demonstrated remarkable success in cancer remission. Therefore, there is a potential benefit of using ICB in PWH to improve the effectiveness of vaccine-induced T-cell responses. Here, we evaluated functional changes in vaccine-induced HIV-specific CD8⁺ T-cell responses by in-vitro PD-1 and TIM-3 blockade during immune stimulation in PWH early ART-treated and vaccinated (Etvac). We compared Etvac to PWH early ART-treated (Et) and chronic ART-treated PWH (Chro). Our results demonstrate a significant increase in the functional responses of vaccine-induced HIV-1-specific CD8⁺ T-cells upon PD-1 but not TIM-3 blockade in cellular proliferation, IFNγ production, and activation. Although no effect was found in Et, similar findings were obtained in HIV-1-specific CD8⁺ T-cells in Chro for combined PD-1/TIM-3 blockade. Moreover, we identify in Etvac a direct association between PD-1 expression in CD8^+^ T cells and in vitro response to anti-PD-1 treatment. Our findings support the potential adjuvant effect of PD-1 blockade to improve HIV-1 therapeutic vaccine efficacy in early ART-treated PWH for future human studies.

## Introduction

Over the last decades, the introduction of antiretroviral treatment (ART) has dramatically improved the control of viral infection and the quality of life in people with HIV-1 (PWH)^1^. However, despite its effectiveness in suppressing viral replication to undetectable levels, ART is unable to eradicate HIV-1, as evidenced by the rapid and consistent viral rebound after treatment interruption^2,3^. The persistence of HIV-1 in the form of a viral reservoir is influenced by complex virus-host interactions and represents a major limitation for cure ^4,5^.

Studies in spontaneous HIV-1 controllers support the critical contribution of HIV-1-specific CD8⁺ T-cell responses to durable viral control in the absence of ART^6,7^. Based on these findings, significant efforts have been directed towards generating potent HIV-1-specific CD8⁺ T-cell responses to achieve a functional cure^8,9^. In this regard, therapeutic T-cell vaccines have been extensively explored^10–13^. However, despite the ability of some therapeutic T-cell vaccine strategies to increase the magnitude and breadth of HIV-1-specific CD8⁺ T-cell responses, their success in the control of viral replication after analytical treatment interruption (ATI) has been limited^13–16^. Therefore, therapeutic T-cell vaccination alone is likely to prove insufficient to control viral replication in vivo, and combinatorial immunoregulatory approaches need to be explored to achieve durable virologic control in the absence of ART^17^.

Inhibitory receptors (IRs), or immune checkpoint molecules, are surface proteins expressed in T-cells and other immune cells that regulate cellular function by dampening TCR signaling^18^. The co-expression of multiple IRs, such as PD-1, TIM-3 or TIGIT, is a feature of T-cell exhaustion in chronic viral infection^18–21^. Additionally, IRs play a relevant role in T-cell activation and are upregulated in vaccine-induced T-cells, which may be limiting their functionallity^18,22^. Several studies have demonstrated the association between the loss of PD-1 expression in CD8⁺ T-cells and improved effector and central memory responses, resulting in an increase of IFNγ, IL-2, higher CD27 and CD62, and better control of viral infections^23,24^. Similar findings have been found through the targeting of TIM-3, leading to stronger virus-specific CD8⁺ T-cell responses in acute viral infection, resulting in the increased production of TNF and IFNγ and higher quality of memory and effector CD8⁺ T-cell responses^25^. Hence, we propose the targeting of IRs in the context of therapeutic T-cell vaccination to enhance vaccine efficacy for HIV-1 cure strategies.

Research in the HIV-1 field has revealed the potential of monoclonal antibodies targeting IRs or immune checkpoint blockade (ICB) in reinvigorating exhausted T-cell responses^26–29^. Moreover, previous studies in chronic SIV infection support the revitalisation of CD8⁺ T-cells by combinatorial use of ICB and vaccines that can induce prolonged viral suppression^29–31^. In addition, the clinical use of ICB for cancer treatment has significantly increased^32^, achieving remarkable outcomes in cancer remission through the reinvigoration of antitumoral immune responses^33^.

Currently, no data on the combinatorial use of ICB and vaccines are available for HIV-1 infection in humans. Here, we assess the effectiveness of ICB in boosting vaccine-induced T-cells to improve vaccine efficacy in PWH. Specifically, we evaluated in vitro the use of anti-PD-1 and anti-TIM-3 antibodies alone or in combination in samples from PWH early ART-treated and vaccinated (Etvac). The Etvac group comprises a unique cohort of early-treated individuals who received a therapeutic vaccine based on an HIVconsv T-cell immunogen after ART initiation, in which vaccine responses demonstrated strong CD8⁺ T-cell HIV-1 inhibitory potential^34,35^. Here, we assessed in vitro functional changes of vaccine-induced HIV-1-specific CD8⁺ T-cell responses regarding cellular proliferation, IFNγ production and activation compared to PWH early ART-treated (Et) and chronic ART-treated PWH (Chro).

Our study revealed that PD-1 blockade significantly enhances functional markers in vaccine-induced HIV-1-specific CD8⁺ T-cells, improving their proliferation and functional activation in the absence of TIM-3 blockade benefit. Notably, we identified distinct cellular clusters in Etvac with higher proliferation, IFNγ production and T-cell activation upon PD-1 blockade, linked to a specific profile of soluble effector markers and antiviral mediators. Additionally, PD-1 expression in CD8⁺ T-cells emerged as a potential biomarker for predicting response to combined therapeutic vaccination and PD-1 blockade. These findings underscore the potential of targeting PD-1 to enhance vaccine-induced CD8⁺ T-cell responses, providing a compelling rationale for further research and clinical exploratory studies.

## Results

### PD-1 blockade during immunogen-specific stimulation enhances the function of vaccine-induced HIV-1 specific-CD8⁺ T-cells

To investigate the effect of ICB on vaccine-induced HIV-1-specific CD8⁺ T-cell responses, we performed an ICB assay in PBMCs from PWH early ART-treated and vaccinated (Etvac, n=13) in the presence of αTIM-3, αPD-1, and αTIM-3+αPD-1 and compared to PBMCs from PWH early ART-treated (Et, n=13) in the control arm of the BCN01 study. Moreover, we included unrelated PBMCs from chronic ART-treated PWH (Chro, n=11) for comparative purposes. The demographic and clinical characteristics of the study groups are detailed in **Table 1**.

**Table 1.**
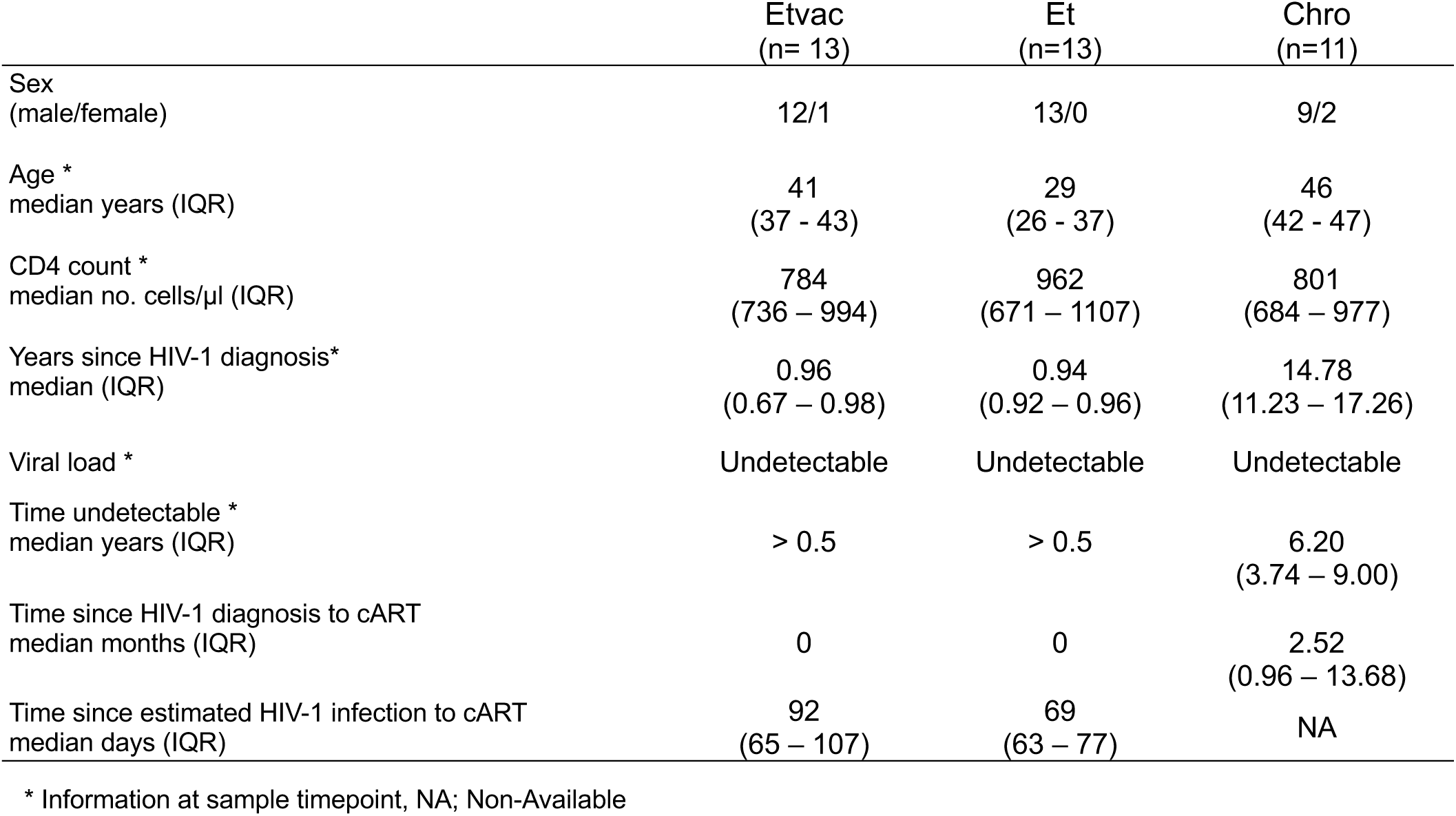
Clinical and demographic characteristics of the study groups.

|  | Etvac<br>(n= 13) | Et<br>(n=13) | Chro<br>(n=11) |
| --- | --- | --- | --- |
| Sex<br>(male/female) | 12/1 | 13/0 | 9/2 |
| Age *<br>median years (IQR) | 41<br>(37 - 43) | 29<br>(26 - 37) | 46<br>(42 - 47) |
| CD4 count *<br>median no. cells/μl (IQR) | 784<br>(736 – 994) | 962<br>(671 – 1107) | 801<br>(684 – 977) |
| Years since HIV-1 diagnosis*<br>median (IQR) | 0.96<br>(0.67 – 0.98) | 0.94<br>(0.92 – 0.96) | 14.78<br>(11.23 – 17.26) |
| Viral load * | Undetectable | Undetectable | Undetectable |
| Time undetectable *<br>median years (IQR) | > 0.5 | > 0.5 | 6.20<br>(3.74 – 9.00) |
| Time since HIV-1 diagnosis to cART<br>median months (IQR) | 0 | 0 | 2.52<br>(0.96 – 13.68) |
| Time since estimated HIV-1 infection to cART<br>median days (IQR) | 92<br>(65 – 107) | 69<br>(63 – 77) | NA |

\* Information at sample timepoint, NA; Non-Available

Briefly, we cultured CFSE-labelled PBMCs with monoclonal antibodies targeting PD-1 and TIM-3 immune checkpoint pathways (αTIM-3, αPD-1 and αTIM-3+αPD-1 or Combo). After seven days in the presence of HIVconsv immunogen-specific peptide pool for Etvac and HIV-1 Gag peptide pool for Et and Chro, we determined the percentage of CFSE^low^ cells, as a readout of cellular proliferation, IFNγ production and HLA-DR⁺CD38⁺ co-expression in CD8^+^ T-cells by flow cytometry (**Fig. 1A-C**). Our results demonstrated a significant increase in proliferation, IFNγ production, and HLA-DR⁺CD38⁺ expression in vaccine induced HIV-1-specific CD8⁺ T-cells in Etvac by αPD-1 (isotype vs. αPD-1; all p<0.05) and Combo (isotype vs. αTIM-3+αPD-1; all p<0.005) but not in αTIM-3 (**Fig. 1D**). No changes in HIV-1-specific CD8⁺ T-cell responses were observed for Et in any of the conditions tested (**Fig. 1E**). By contrast, αPD-1 significantly increased the proliferative capacity, IFNγ production, and HLA-DR⁺CD38⁺ expression of HIV-1-specific CD8⁺ T-cells in Chro (isotype vs. αPD-1; all p<0.05) (**Fig. 1F**). Interestingly, αTIM-3 alone increased the proliferation of HIV-1-specific CD8⁺ T-cells (isotype vs. αTIM-3; p = 0.03) and Combo condition increased all functional markers (isotype vs. αTIM-3+αPD-1; all p<0.05) (**Fig. 1F**). These results indicate that αPD-1 alone or in combination with αTIM-3 enhances the function of vaccine-induced HIV-1-specific CD8⁺ T-cell responses in Etvac and reinvigorates HIV-1-specific CD8⁺ T-cell responses in Chro during antigenic stimulation.

**Figure 1.**
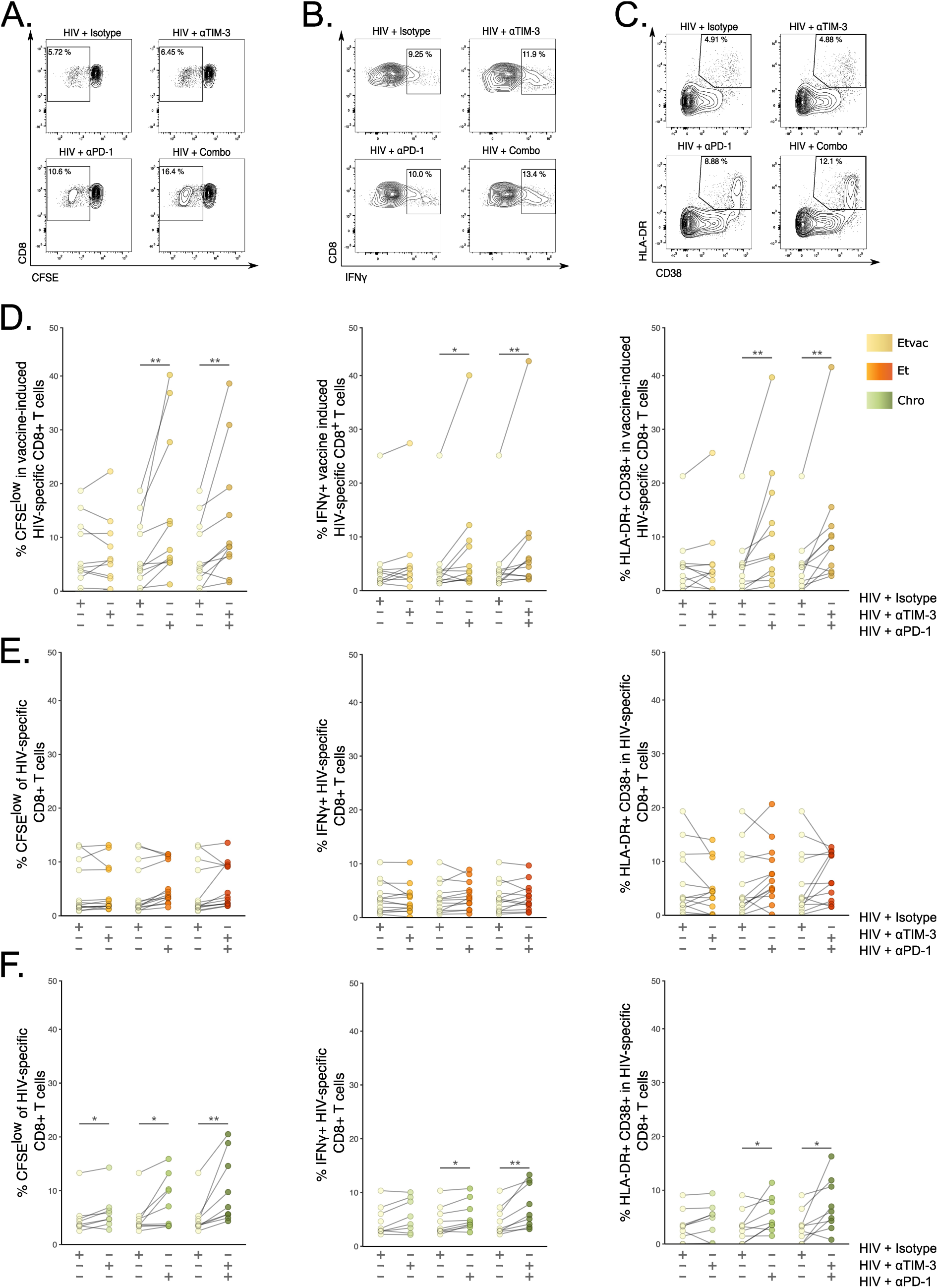
Functional changes in HIV-1-specific CD8⁺ T cells. Representative flow cytometry plots gated on CD8^+^ T-cells for a Chro individual showing A. Cellular proliferation (CFSElow), B. IFNγ production, and **C**. HLA-DR⁺CD38⁺ expression. Pair-wise comparison on the percentage of CFSE^low^, IFNγ and HLA-DR⁺CD38⁺ expression in response to the corresponding HIV peptides (HIV-1-Gag peptide pool for Et and Chro groups and HIVConsv peptide pool for Etvac) in the presence of αPD-1, αTIM-3, αPD-1+αTIM-3 (Combo) antibodies and isotype control antibodies in **D.** Etvac, **E.** Et and **F.** Chro study groups. The dots represent a sample tested. The Wilcoxon matched-pairs signed ranked test was used to calculate statistical differences. P-values: ns > 0.05, *<0.05, **<0.005 and ***<0.0005.

To further evaluate the potential contribution of αTIM-3 in the Combo treatment (αTIM-3+αPD-1), we transformed all the functional data into Fold-change (FCh) values relative to the isotype control and performed a linear mixed-effects model using αPD-1 as the comparator. As shown in the waterfall plots of **Fig. 2A**, we detected a significant increase in FCh proliferation, IFNγ production, and HLA-DR⁺CD38⁺ (all, p<0.00005) in HIV+αPD-1 vs. HIV, consistent with the previous analysis. However, no significant differences were found between HIV+αPD-1 and HIV+Combo for any functional markers (**Fig. 2A)**. Similar results were obtained when analysing Etvac and Et independently (**Fig. 2B-C**). Hence, our data indicate an independent effect of αPD-1 as the driver of the changes observed in vaccine-induced HIV-1-specific CD8⁺ T-cells in Etvac with no additional benefit of αTIM-3 in the Combo. Nevertheless, we observed an increase in proliferation (p = 0.038) and IFNγ production (p = 0.033) in the absence of changes in HLADR^+^CD38^+^ in Chro between HIV+αPD-1 and HIV+Combo condition (**Fig. 2D**), implying a specific effect of Combo treatment in Chro. Therefore, our findings indicate a unique contribution of αTIM-3 in the recovery of exhausted HIV-1-specific CD8⁺ T-cells in combination with αPD-1 in these participants who initiated ART treatment in chronic stages of the infection.

**Figure 2.**
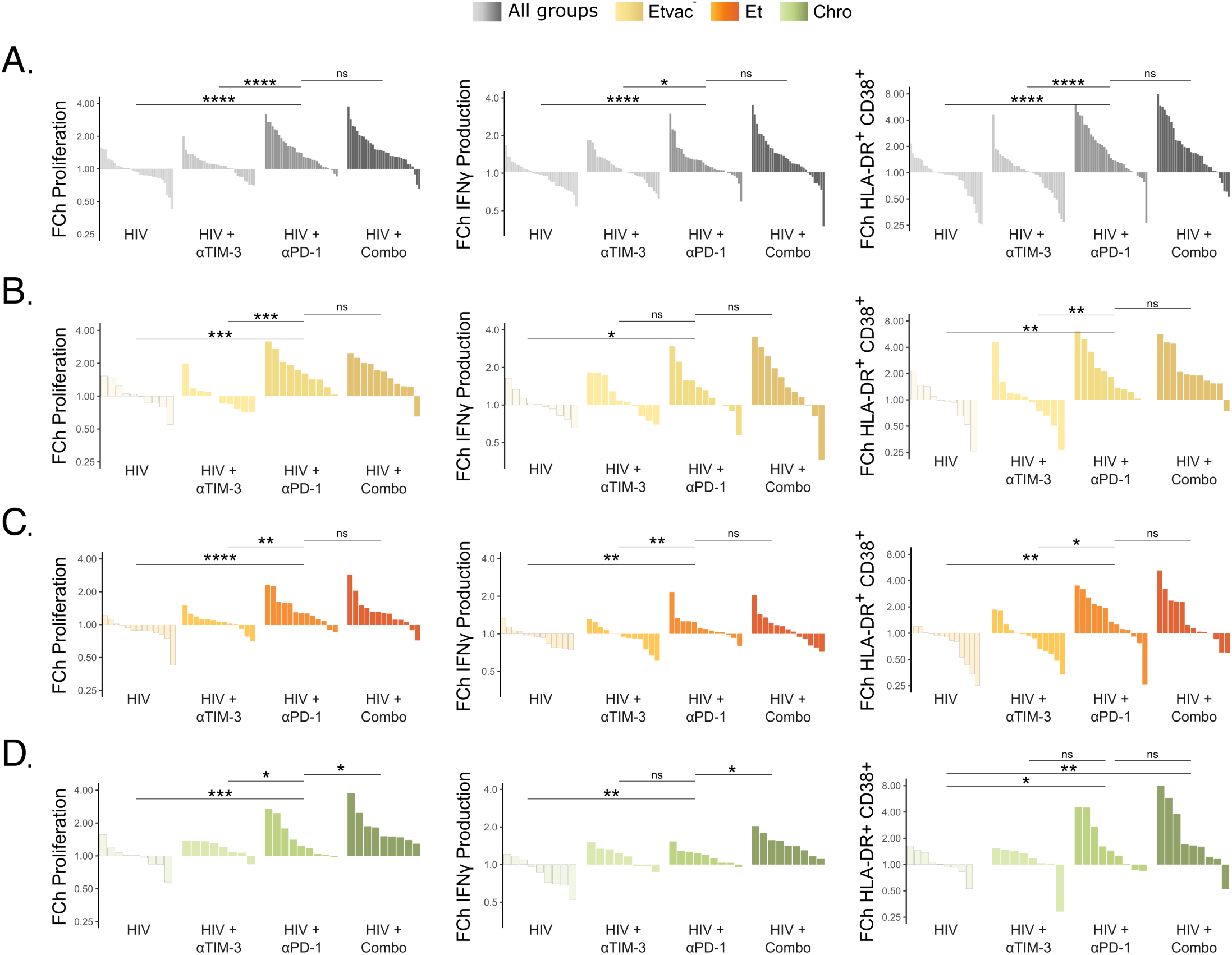
Waterfall plots of functional changes in HIV-1-specific CD8⁺ T cells in Fold Change (FCh). Waterfall plots represent the magnitude of proliferation (CFSE^low^), IFNγ and HLA-DR⁺CD38⁺ as FCh of each condition from the Isotype control. **A.** All study groups combined (n=33), **B.** Etvac (n=11), **C.** Et (n=13) and **D.** Chro (n=9). The bars represent the FCh per sample tested. The statistical linear mixed-effects model calculates differences using αPD-1 as a reference group. P-values: ns>0.05, *<0.05, **<0.005 and ***<0.0005.

### PD-1 blockade during antigenic stimulation elicits vaccine-induced HIV-1 specific-CD8⁺ T-cell clusters co-expressing T-cell functional markers

To characterize the effect of ICB in CD8^+^ T-cell responses, we conducted a correlation matrix analysis across functional markers tested in the presence of αTIM-3, αPD-1 and Combo. For Etvac, we observed a positive correlation between proliferation and activation in the presence of HIV-1 peptides (ρ=0.93; p < 0.0005) and HIV+αTIM-3 (ρ=0.77, p < 0.05). Moreover, we observed a significant positive correlation between all functional markers in αPD-1 treatment (ρ>0.85; p < 0.005), while only proliferation and IFNγ production (ρ=0.71; p < 0.05) were statistically significant in Combo treatment. Finally, we only detected a significant correlation between proliferation and activation (ρ=0.64; p < 0.05) in Combo for Et in the absence of any correlation for Chro (**Fig. 3**).

**Figure 3.**
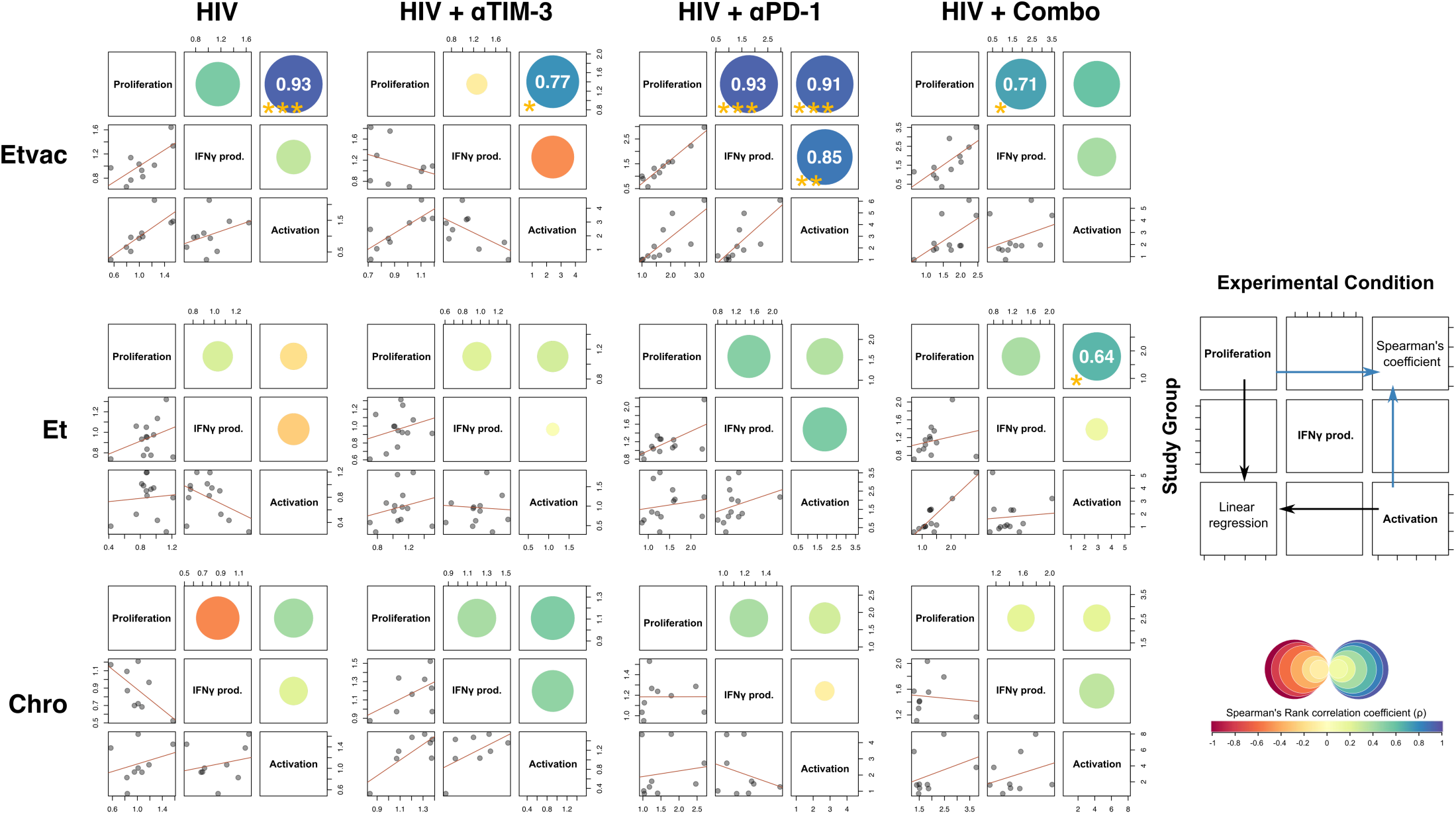
Correlations between functional markers of HIV-1-specific CD8⁺ T-cell responses. Correlograms indicate correlations between parameters across study groups: Etvac (n=11), Et (n=13) and Chro (n=9) for each experimental condition (HIV-1 for peptide stimulation, HIV+αTIM-3, HIV+αPD-1 and HIV+Combo). The top of the diagonal shows Spearman’s rank correlation coefficient with p-value<0.05. The colour and size of the circles indicate the strength and direction of the correlation. The bottom of the diagonal represents the linear regression. p-values: ns>0.05, *<0.05, **<0.005 and ***<0.0005.

Next, to further validate the association across functional parameters observed in correlation analysis of vaccine-induced HIV-1-specific CD8⁺ T-cells, we performed a net-SNE dimensionality reduction and unsupervised cell clustering through KNN algorithms to analyze 127,142 CD8^+^ cells from Etvac under both, the isotype and αPD-1 conditions. We defined 11 cellular clusters distributed according to the relative intensity of marker expression of 4 parameters, as shown in **Fig. 4A-B**. Remarkably, out of the 11 clusters identified, clusters #1 and #6 showed higher intensities in HLA-DR, CD38 and IFNγ, concomitant with a reduction of CFSE intensity (cell proliferation) (**Fig. 4C**). In addition, cluster #1 exhibited higher intensity for IFNγ compared to #6. Moreover, a paired comparison of the cell frequencies in each cluster between isotype and αPD-1 demonstrated that cluster #1 was predominantly comprised of a single Etvac cell in αPD-1 treatment; meanwhile, cluster #6 composition was significantly increased in 8 Etvac in αPD-1 treatment compared to isotype control (**Fig. 4D**). Since the use of αPD-1 has been proposed to induce reactivation of HIV-1 latently infected CD4⁺ T-cells^28^, we performed ultrasensitive p24 in the supernatants from ICB experiments in Etvac. Interestingly, we detected p24 values above the limit of quantification (LOQ) under all experimental conditions, including superantigen stimulation and unstimulated conditions. Remarkably, the median levels of p24 were lower in the αPD-1 condition in Etvac samples below the LOQ (median= 0.018 pg/mL, IQR 0.023-0.016) (**Fig. 4E**).

**Figure 4.**
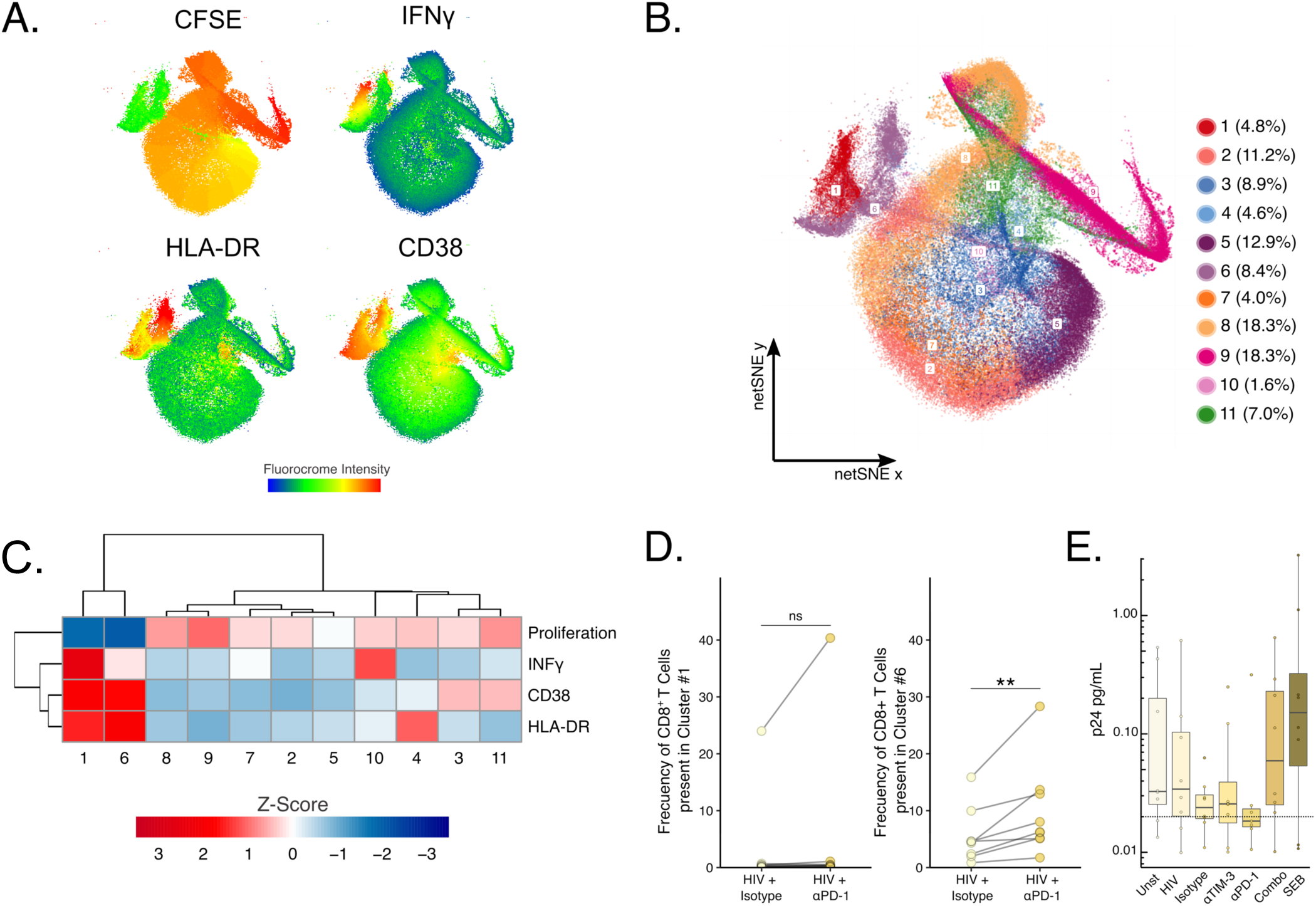
Unsupervised analysis of vaccine-induced HIV-1-specific CD8⁺ T-cells in αPD-1 treatment. **A.** netSNE visualization of flow cytometry data for Isotype and PD-1 blockade condition in Etvac (127.142 events). Each panel represents the fluorochrome intensity of CFSE, IFNγ, HLA-DR and CD38 for each recorded event. **B.** Clustering analysis of cells in netSNE space by KNN algorithm. The percentage of the total cells comprised in each cluster is indicated. **C.** Heatmap of the median normalised biexponential-transformed intensity for each marker. The dendrogram shows the association between clusters based on the intensity of each fluorochrome. Two major nodes indicate proliferating (left node, #1, #6) and no proliferating (right nodes). **D.** Pairwise comparison of cluster #1 and #6 composition indicates the frequency of CD8⁺ T cells inside the cluster in isotype and αPD-1 conditions. **E.** Ultrasensitive p24 determination in the supernatant seven days after cell culture. The dotted line represents the limit of quantification (LOQ). P value<0.005(**). Unst (Unstimulated) SEB (Staphylococcal enterotoxin B). The Wilcoxon matched pairs signed ranked test was used to determine statistical significance. p-values: ns>0.05, *<0.05, **<0.005 and ***<0.0005.

These data indicate that the functional markers evaluated are enhanced proportionally and concurrently by αPD-1 in Etvac. Moreover, our findings provide evidence for specific vaccine-induced cellular clusters elicited by αPD-1 in Etvac co-expressing functional markers and potentially associated with a reduction of latently infected cells (p24 levels). These data suggest at least some level of reservoir reactivation by αPD-1, which may facilitate the clearance of p24-expressing cells by vaccine-induced HIV-1-specific CD8⁺ T-cell responses.

### PD-1 blockade during antigenic stimulation generates a 17-cytokine signature profile of antiviral factors and degranulation molecules

Next, to identify a potential immune signature associated with enhancing vaccine-induced responses in Etvac by αPD-1 treatment, we conducted a 17-cytokines multiplex assay in supernatants from the ICB experiments. Using linear mixed-effects model analysis, we identified a signature of 13 soluble factors in response to αPD-1 across study groups, including MIP-1α, Granzyme-B, IFNγ, IL-13, IL-5, sCD137, Perforin, sFasL, MIP-1β, GM-CSF, Granzyme-A, IL-10, and IL-2 in most of the samples exposed to HIV-1 antigenic stimulation and αPD-1 or Combo treatment (**Fig. 5A**, light and dark brown). This signature includes relevant chemokines and antiviral HIV-1 factors, such as MIP-1α and MIP-1β, as well as effector cytokines and degranulation molecules, such as Perforin, Granzyme-A or Granzyme-B necessary for the elimination of infected cells. Additionally, we used STRING tool^36^ to perform a functional Enrichment Analysis on the 13-set of soluble factors differentially expressed in response to αPD-1. The analyses revealed several biological processes, including immune responses, cytokine-mediated signalling, response to other organisms, and defence response (**Fig. S2A**). The enrichment analysis also revealed KEGG pathways associated with various autoimmune diseases, in addition to T-cell signalling (hsa04660) and NK-mediated cytotoxicity pathways (hsa04650) (**Fig. S2B**). Nonetheless, linear mixed-effects model intra-group analysis revealed significant differences in response to αPD-1 and antigenic stimulation. Particularly, Etvac individuals produced more IL-13, MIP1b, and Perforin, while Chro produced more IL-10 and IL-2 (**Fig. 5B**). Meanwhile, the intergroup analysis indicated an Etvac-specific signature of cytokines in response to αPD-1 characterised by increased IL-5 and IL-13 production (**Fig. 5C**). These data suggest that, during HIV-1 antigenic stimulation, PD-1 blockade can induce a specific cytokine signature, which may enhance anti-viral vaccine-induced immune responses, particularly in Etvac individuals, while generating homeostatic and anti-inflammatory immune responses in Chro individuals.

**Figure 5.**
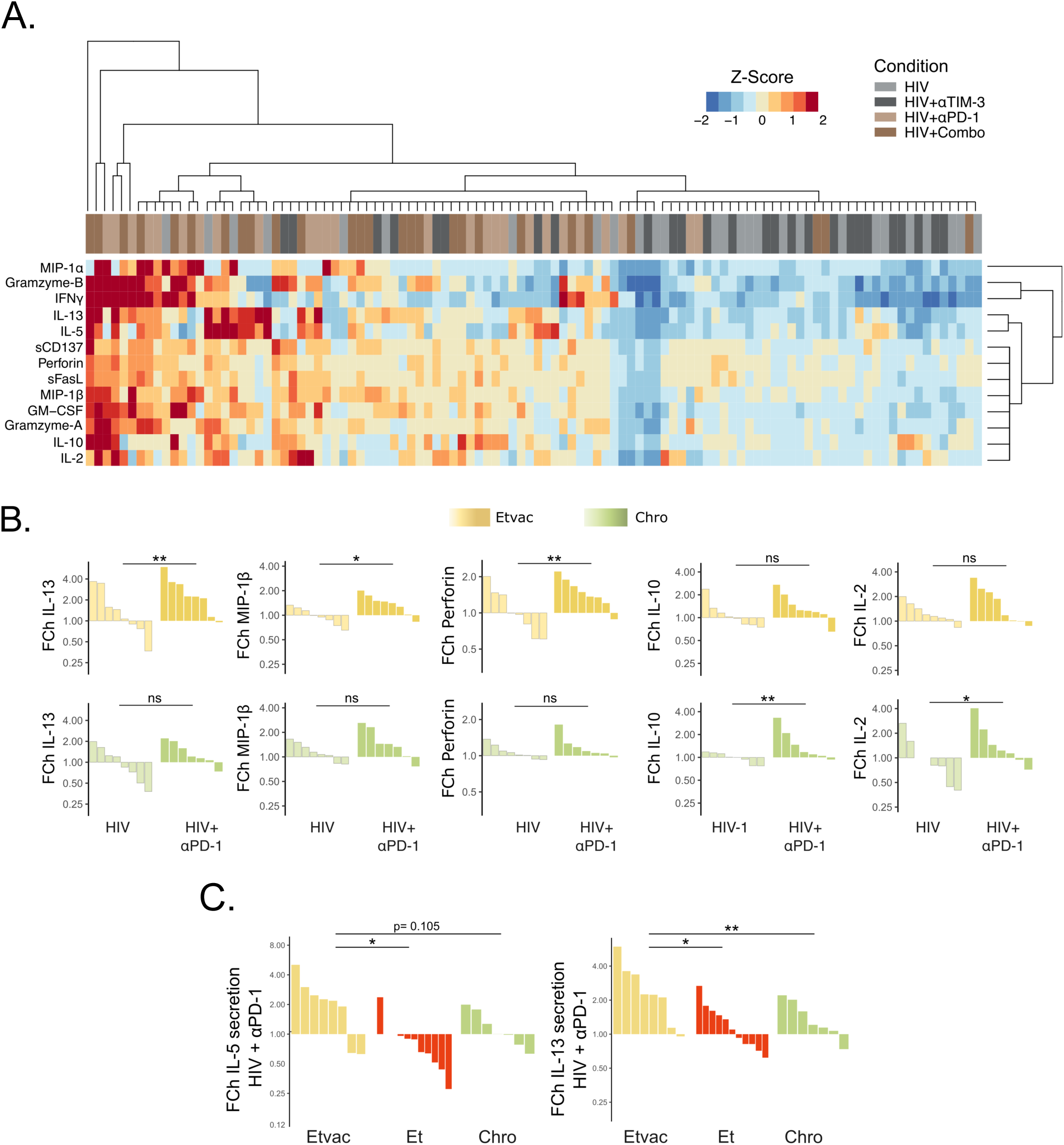
Cytokine profile in response to αPD-1. **A.** Heatmap of the Cytokine profiling across study groups (n=104 experimental conditions). **B.** Waterfall plots show secreted cytokines in Etvac (n=8) or Chro (n=7) in response to αPD-1 represented as the fold Change (FCh) from the isotype control condition. **C**. Waterfall plots of IL-5 and IL-13 production Etvac (yellow), Et (red) and Chro (green) groups upon αPD-1 treatment. The heatmap colour scale indicates the standardized FCh of each cytokine across all samples. The bars represent a sample FCh per individual tested. The linear mixed model was used to determine statistical significance using αPD-1 as a reference group. p-values: ns>0.05, *<0.05, **<0.005 and ***<0.0005.

### PD-1 expression levels in CD8⁺ T cells correlate with in vitro functional responsiveness to blockade in Etvac during antigenic stimulation

To evaluate potential markers of responsiveness to in vitro ICB, we performed PD-1, TIM-3, HLA-DR, CD38 and CD8 staining in cryopreserved PBMCs before ICB studies. No differences in TIM-3 and PD-1 expression frequency in CD8⁺ T cells were observed among study groups (**Fig. 6A**). Nonetheless, we observed two subgroups of Etvac, with a median of 30.7% (n=5) and a 12.3% (n=5) of PD-1 expression in CD8 T-cells. Additionally, we found a significantly higher expression of HLA-DR⁺CD38⁺ in CD8^+^ T cells from Etvac compared to Et and Chro (**Fig. 6B**). Moreover, CD38 expression was significantly lower in Chro than in Etvac and Et (**Fig. 6B**). Additionally, correlation analysis demonstrates that basal PD-1 expression in total CD8^+^ T cells correlated with proliferation (R=0.77, ρ=0.014) and HLA-DR⁺CD38⁺ (R= 0.67 ρ=0.039) in the presence of αPD-1 for Etvac (**Fig. 6C**). These findings were exclusive for Etvac in the absence of correlations for any other groups (**Fig. 6C-D**). Thus, according to our findings, baseline levels of PD-1 expression in CD8⁺ T cells are associated with the increase of vaccine-induced functional CD8⁺ T cell responses in the presence of αPD-1 during HIV-1 antigenic stimulation in Etvac. These data support the use of PD-1 expression in CD8 T cells as a potential biomarker of response to αPD-1 treatment when considering its use in combination with therapeutic T-cell vaccination.

**Figure 6.**
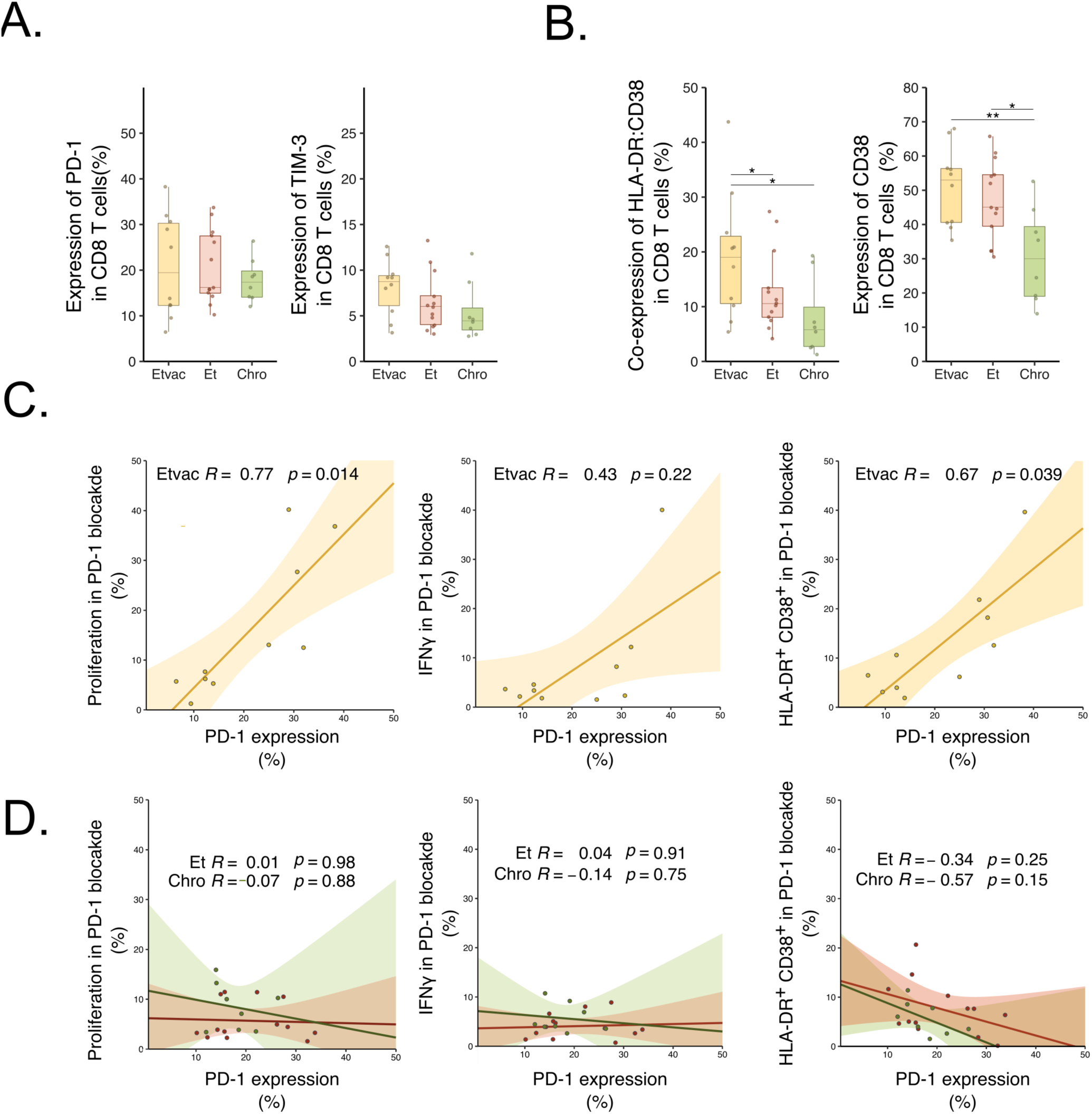
Functional responses are associated with the expression of PD-1 in CD8⁺ T cells. **A** Basal expression of inhibitory receptors and HLA-DR⁺CD38⁺ expression in PBMCs from Etvac (n=10), Et (n=13) and Chro (n=9) individuals before the immune checkpoint blockade assay. **B** Correlations between the percentage of CFSE-Low, IFNγ production and HLA-DR⁺CD38⁺ in anti-PD-1 blockade and PD-1 expression in CD8^+^ T cells. Mann–Whitney test was used to compare differences between groups. Spearman correlations between Basal PD-1 levels (frequency) and the functional parameters determined on day 7 upon PD-1 blockade, p-values: ns>0.05, *<0.05, **<0.005 and ***<0.0005.

## Discussion

Therapeutic T-cell vaccination is considered an essential component in achieving an effective HIV-1 cure^9,17^. However, the limited control of HIV-1 infection observed in vaccine trials after an ATI suggests that vaccine-induced HIV-1-specific responses are constrained by several factors, including the expression of IRs following vaccine-induced immune activation^18^. Immune interventions based on ICB have been proposed to tackle T-cell exhaustion in HIV-1 chronic infection models and cancer ^37^. However, single ICB interventions in HIV-1 preclinical studies have shown to be insufficient for eliminating the viral reservoir, even though they can partially reinvigorate pre-existing virus-specific T-cell responses^38^. Thus, there is an urgent need to explore the potential of using a combination of T-cell vaccines and ICB to achieve the HIV-1 cure^17^.

Currently, no data are available on the combined use of these immunotherapeutic approaches in PWH. Existing data only explored therapeutic T-cell vaccines in combination with PD-1/PD-L1 blockade in preclinical models of chronic viral infection, such as chronic SIV infection in NHP^30,31^. Notably, studies on the chronic SIV/rhesus macaque model suggest that αPD-1 treatment could enhance the cytolytic potential of vaccine-induced CD8⁺ T-cell responses during T-cell exhaustion^29,31^. This was evidenced by the increased frequency of CD8⁺ T-producing perforin and granzyme B cells in blood and lymph nodes. This, in turn, led to improved localisation of these cells in the germinal centres of B cell follicles, ultimately leading to a decrease in the viral reservoir when a combinatorial vaccination strategy (CD40L plus TLR7 agonist– adjuvanted DNA/MVA vaccine and αPD-1 treatment) was employed in chronic SIV infection^31^.

Our study evaluates the impact of combining ICB during vaccine immune activation in samples from individuals who receive early ART after recent HIV-1 acquisition (Etvac) in the context of the BCN01 vaccine clinical trial^35^. Importantly, Etvac received a prime-boost vaccine regime that induced a selective expansion of T-cell responses toward HIVconsv immunogen that shifted from less than a median of 2% to 48% of total HIV-1 responses at the peak of immunogenicity^35^. By using PBMCs from Etvac at the immunogenicity peak and comparing CD8⁺ T-cell responses to unvaccinated early-treated individuals from the control arm of the BCN01 clinical trial, we assessed the in vitro effect of αPD-1 and αTIM-3 alone or combined on the function of potentially de-novo-induced HIV-1-specific CD8⁺ T-cell responses that were particularly subdominant before vaccination^35^

Our investigation revealed that αPD-1 enhanced the functional markers in vaccine-induced HIV-1-specific CD8⁺ T-cells during HIV-1 antigenic stimulation independently of αTIM-3. Moreover, we validated the concurrent enhancement of functional parameters upon αPD-1 in Etvac, demonstrating the expansion of vaccine-induced CD8⁺ T-cells producing IFNγ and becoming activated (by co-expressing HLA-DR and CD38) in the presence of vaccine-immunogen specific peptides and αPD-1 treatment. Remarkably, an Etvac individual who was later defined as an ATI controller in the BCN02 clinical trial^39^ was identified in the Etvac cellular cluster #1, characterized by higher production of IFNγ, HLA-DR, and CD38 compared to cluster #6. Conversely, PD-1 blockade did not significantly enhance existing HIV-1-specific CD8⁺ T-cell responses in Et, suggesting that vaccination had led to an expansion of immunogen-specific cells that were able to profit from αPD-1 treatment. These findings highlight the specific effect of αPD-1 during immune activation in boosting the HIVconsv vaccine-induced HIV-1-specific CD8⁺ T-cell responses, which can be attributed to the expression of PD-1 induced by vaccination.

These data overcome limitations of previous studies using the immunodominant Gag protein as immunogen for vaccination in chronic infection models, where T-cell responses were measured with a Gag peptide stimulation, limiting the assessment of the de-novo-induced HIV-1-specific CD8⁺ T-cell responses^29–31^. Interestingly, we observed a combined effect of αPD-1 and αTIM-3 in reinvigorating HIV-1-specific CD8⁺ T-cell responses in Chro, aligning with reports of ICB facilitating the recovery of exhausted T cells in chronic HIV-1 infection^25,38^. Nonetheless, complementary information is required to fully understand the contribution of combinatorial TIM-3 and PD-1 blockade during stimulation in restoring T-cell function in HIV-1 chronic infection as these may comprise different T cell subsets than those present among the vaccine-induced T cell population in Etvac.

Additionally, the use of ICB antibodies targeting PD-1, CTLA-4, and other immune checkpoints showed promise in perturbing HIV-1 latency during ART^25,32,34,35^,. Ideally, ICB antibodies could be able to reverse HIV-1 latency and boost HIV-1-specific immunity, generating a "shock and kill" strategy with a single therapeutic agent^28,40^. In this study, we evaluated changes in HIV-1 reactivation by ultrasensitive p24 detection in supernatants in the presence of HIV-1 peptides and ICB antibodies. We found that αPD-1 decreased p24 levels below the limit of quantification in Etvac compared to control conditions. These results align with the possibility that the enhancement of vaccine-induced HIV-1-specific CD8⁺ T-cell responses following PD-1 blockade was responsible for the decreased HIV-1 reactivation, likely due to the enhanced killing of reactivated CD4⁺ T-cells displaying viral antigens that were poised to react by PD-1 blockade, directly supporting combination strategies. However, limitations in the number of cells available prevented an accurate evaluation of the levels of proviral HIV-1 DNA or performing Q-VOA assay in our study. Additional experiments are needed to explore the potential of combinatorial T-cell vaccination and αPD-1 to reduce the viral reservoir.

We identified a signature of 13 cytokines or soluble factors in response to HIV-1 peptide stimulation during αPD-1 treatment, alone or combined with αTIM-3, across study groups. For Etvac, the profile was characterised by the secretion of Perforin, MIP-1β, IL-5 and IL-13 while Chro produced IL-10 and IL-2. This observation indicates an increased effector and anti-viral responses in Etvac while suggesting a shift to homeostatic and anti-inflammatory T-cell responses in Chro individuals. Previous studies using the HIVconsv vaccine demonstrated similar cytokine profiles characterised by the production of MIP-1α, MIP-1β, Granzyme-B, Granzyme-A, IFNγ, and IL-2, which persist for up to 2 years in vaccine-elicited human T cells after vaccination^41^. However, αPD-1 treatment in Etvac further enhances these responses and increases the cytotoxic potential of CD8^+^ T cells through the secretion of perforin, IL-5 and IL-13 production. On the other hand, the shift in Chro αPD-1 signature, driven by IL-10, may have adverse effects on T-cell effector function and potentially result in immunosenescence favouring HIV-1 persistence^42^. Consequently, further research is needed to understand the signature profile of αPD-1 treatment during chronic HIV-1 infection and address the mechanism behind it.

Over the last years, multiple clinical trials have assessed the safety and tolerability of ICB therapeutic strategies in cancer patients with HIV-1, increasing the data available that support the feasibility of ICB treatment in PWH^43–47^. Nonetheless, identifying biomarkers of response to ICB is essential for the personalized use of these therapeutic immune interventions in PWH. In our study, the expression of PD-1 in total CD8⁺ T-cells was consistent with previous reports in vaccinated individuals from the BCN01 clinical trial^35^ Our data suggests that the frequency of PD-1 expression in total CD8^+^ T cells at the peak of vaccine-induced immunogenicity was associated with response to PD-1 blockade in terms of increased proliferation and T-cell activation. These findings suggest that PD-1 expression in CD8⁺ T-cells could be further investigated as a biomarker to stratified PWH for combined therapeutic T-cell vaccination and αPD-1 treatment. Interestingly, these results contrast with the lack of predictive value of PD-1 expression observed in Chro, which is consistent with observations in the cancer field where PD-1 expression is not considered an excluding or definitive predictive factor for patient stratification in immunotherapies^48^.

Collectively, our findings support the role of PD-1 beyond T-cell exhaustion^18^ as a promising target to enhance vaccine-induced CD8^+^ T-cell responses, providing an in vitro proof of concept for using αPD-1 treatment in combination with therapeutic T-cell vaccines to improve vaccine efficacy. PD-1 blockade may alleviate the functional constraints imposed by the PD-1^+^ effector vaccine-induced CD8^+^ T-cells upon immune activation, allowing the targeting of the immune privilege site where the HIV-1 persistence is present^49,50^. Current research in combinatorial immunoregulatory approaches, such as the combined use of therapeutic T-cell vaccines and TLR7 agonists, is currently being tested in different clinical trials with encouraging results, including AELIX-003 or NCT04357821^51–54^.

Our study offers novel insights into developing combinatorial immune interventions aimed at enhancing cellular immunity through the integration of therapeutic T-cell vaccines with PD-1 immunomodulatory molecules. Furthermore, exploring PD-1 expression in CD8^+^ T-cells at the peak of immunogenicity as a predictive biomarker could significantly enhance patient stratification, thereby paving the way for more effective, personalized immunotherapies and impacting the pursuit of an HIV-1 cure.

## Methods

### Study groups

The study groups were selected from the BCN01 trial (NCT01712425). The BCN01 trial was a phase I, open-label, non-randomised, multicentre therapeutic vaccination study to evaluate the safety and immunogenicity of a vaccination regimen of prime/boost with ChAdV63.HIVconsv/MVA.HIVconsv in PWH in early ART treatment^35,55^. From the study, we selected cryopreserved PBMCs from PWH early treated and vaccinated (Etvac; n=13) at the immunological peak of vaccine response and cryopreserved PBMCs at the equivalent time point from the study control arm, PWH early treated but not vaccinated (Et; n=13). In addition, we include PWH with at least six years of suppressive ART initiated during the chronic infection phase (Chro; n=11). The clinical and epidemiological characteristics of the study groups are summarised in **Table 1**.

### Ethics Statement

All subjects provided their written informed consent for the purpose of research. The project was approved by the institutional review board of Germans Trias i Pujol Hospital (PI-17-039). The study was conducted under the principles expressed in the Declaration of Helsinki.

### PD-1, TIM-3 and PD-1+TIM-3 checkpoint blockade assay

We selected cryopreserved PBMCs from Etvac (n=11), Et (n=13) and Chro (n=9) as previously mentioned. PBMCS were thawed and resuspended in R10 media (RPMI media containing 10% FCS, 100 U/mL Penicillin and 0.1 mg/mL Streptomycin) and rested at 37 °C with 5% CO_2_ for 4 hours. Next, to evaluate cellular proliferation, PBMCs were labelled with Vybrant Carboxyfluorescenin diacetate succinimidyl ester (CFSE) dye (Vybrant™ CFDA SE Cell Tracer, Invitrogen) before adding blocking antibodies as previously described^26^. Briefly, 10^6^ PBMCs were incubated with 0.5 μM CFSE dye for 5 minutes at room temperature. After incubation, CFSE-labelled PBMCs were extensively washed with 1x PBS and incubated for 20 minutes at 37 °C to remove the excess CFSE dye. Then, CFSE-labelled PBMCs were washed with 1x PBS and rested overnight at 37 °C. The next day, CFSE-labelled cells were incubated under R10 complemented medium with 1 μg/ml of anti-CD28/CD49d (BD, Pharmigen) in the following conditions: 1) unstimulated, 2) Staphylococcal enterotoxin B (SEB, 1 μg/ml, Sigma-Aldrich), 3) HIV-1-Gag peptide pool (2 μg pool peptide/ml) for the non-HIVCons vaccinated Et and Chro groups, 4) HIVCons vaccine-immunogen specific peptide pools (2 μg pool peptide/ml) for Etvac. All these conditions were tested in the absence or presence of antibodies targeting PD-1, TIM-3 and TIM-3+PD-1 together with the isotype control antibodies. For PD-1 blockade (αPD-1), we included anti-human PD-1 IgG4 S228P antibody (10μg/ml, 1B8/HuPD1B-3, Merck & Co., Inc., Rahway, NJ, USA). For TIM-3 blockade (αTIM-3), we used anti-human TIM-3 IgG4 antibody (5 μg/ml, Merck & Co., Inc., Rahway, NJ, USA). Finally, we included αTIM-3+ αPD-1 and used anti-human RSV-IgG4 as an isotype control antibody (5 μg/mL, 60AGK S228P, Merck & Co., Inc., Rahway, NJ, USA). On day six, CFSE-labelled cells were boosted with HIV-1-Gag or vaccine-specific peptide pools in the presence of Brefeldin A solution (BD, Biosciences) and Monensin solution (BD Biosciences). The next day, CFSE-labelled cells were surface and intracellularly stained with a panel of antibodies, as described in the section below. The culture supernatants were collected for multiplex and p24 ultrasensitive analyses.

### Polychromatic flow cytometry

Cryopreserved PBMCs were thawed and rested in R10 for 4 hours at 37 °C and 5% CO_2_. Then, we performed a basal immune phenotype and stained 10^6^ PBMCs for 25 minutes with the Live/Dead probe (LIVE/DEAD™ Near-IR Dead Cell Stain, Invitrogen) at room temperature to discriminate dead cells. Cells were washed with 1x PBS and surface stained with CD8 (AmCyan, clone RPA-T8, BD), HLA-DR (PerCP-Cy5.5, clone G46-6, BD), CD38 (APC, clone HB-7, Biolegend), PD-1 (BV421, clone EH12.1, BD) and TIM-3 (PE, clone F38-2E2, Biolegend) antibodies for 25 minutes. Then, samples were washed with 1X PBS and fixed in 1% formaldehyde before acquisition on a FACSCanto™ II (Beckmann Coulter) using the FACS DiVa software (BD, Biosciences). Also, we performed an immune phenotype for checkpoint blockade experiments to monitor cellular proliferation by CFSE loss and cellular function by IFNγ production and HLA-DR^+^CD38^+^ co-expression. Briefly, PBMCs were labelled with a Live/Dead probe (LIVE/DEAD™ Near-IR Dead Cell Stain, Invitrogen) and then stained with the surface markers CD8⁺ (AmCyan), HLA-DR (PerCP Cy5.5), PD-1 (BV421), CD38 (APC) and TIM-3 (PE). Subsequently, cells were washed, fixed and permeabilised with a Fix&Perm kit (Thermo Fisher Scientific) for intracellular cytokine staining with IFNγ (Pe Cy7). Stained samples were fixed in 1% formaldehyde and acquired on FACSCanto™ II (Beckmann Coulter) using the FACS DiVa software (BD, Biosciences). Data were analysed using FlowJo v10.6 (Tree Star Inc). To determine the percentage of HIV-1-specific CD8⁺ T-cell responses, we subtracted the percentage of CFSE lost, IFNγ production, and HLA-DR⁺CD38⁺ expression from the unstimulated background to the HIV-1 peptide stimulation (**Fig. S1**). We performed two technical replicates for HIV-1-specific CD8⁺ T-cell responses. We considered a response positive after background subtraction (mean of two technical replicates) used as the cut-off value. We recorded a median of 7.000 CD8⁺ T-cell events for each independent sample.

### Multiplex cytokine assay and ultrasensitive p24 assay

The MILLIPLEX MAP Human CD8⁺ T-Cell Magnetic Bead Panel (Merck Millipore Darmstadt, Germany) was used to quantify GM-CSF, sCD137, IFNγ, sFas, sFasL, Granzyme A, Granzyme B, IL-2, IL-4, IL-5, IL-6, IL-10, IL-13, MIP-1α, MIP-1β, TNF and Perforin in two technical duplicates from cell culture supernatants at day seven. Following the manufacturer’s instructions, plates were read by Luminex 200, Austin Luminex, USA. Data were analysed using MILLIPLEX Analyst 5.1 software (Merck Millipore Darmstadt, Germany). In addition, we used cell culture supernatants to quantify p24 levels using an ultrasensitive determination HIV p24 digital immunoassay. For the assay, cell culture supernatants were harvested by centrifugation and inactivated by mixing equal supernatant and inactivation buffer (1% Triton x-100 in 0.5% Casein buffer and 50% Hi-FBS) for 15 minutes. The p24 was quantified in technical duplicates as previously described^56,57^. Values below 0.01 pg/mL are considered background, the limit of quantification (LOQ) is 0.02 pg/mL.

### Unsupervised immunophenotype data analysis

The phenotypic and functional characterization of cellular populations was analysed using t-Distributed Stochastic Neighbor Embedding (t-SNE)^58^ and net-SNE^59^ dimensionality reduction algorithms to visualize single-cell distributions in two-dimensional maps. Briefly, compensated fluorochrome intensities of total CD8⁺ live cells were z-normalised, and a randomly selected subset of cells, at least 1,000 cells per sample, was passed through the t-SNE algorithm. The resulting t-SNE dimension was then used to predict the position of all remaining CD8⁺ T cells acquired per sample from each group using the net-SNE algorithm based on neural networks. In parallel, we discovered cell communities using the Phenograph clustering technique. It operates by computing the Jaccard coefficient between nearest neighbours, which was set to 30 in all executions, and then locating cell communities (or clusters) using the Louvain method^60,61^. The method creates a network indicating phenotypic similarities between cells. The netSNE maps included representations of the identified cell communities. Additionally, we built a heatmap with the clusters in the columns and the markers of interest in the rows to better comprehend the phenotypical interpretation of each cluster. The colour scale displays each marker’s median intensity on a biexponential scale. We calculated quantitative assessments of cellular clusters in the percentage of cells for each sample to analyse and compare the distribution between HIV+Isotype and HIV+αPD-1 stimulation, similar to the supervised flow cytometry analysis.

### Statistics

Descriptive statistics of centrality and dispersion are reported as median and interquartile range, respectively. Bivariate analysis was conducted using nonparametric methods: Wilcoxon matched-pairs signed rank test for paired median changes between conditions and Spearman linear correlation coefficient to study the association between continuous variables. All statistics were programmed and performed using the R statistical package^62^. Data including frequency of proliferation, IFNγ, CD38/HLA-DR during HIV stimulation were transformed into Fold Change values (FCh) as the ratio between each experimental condition (αTIM-3, αPD-1 and αTIM-3+αPD-1) and the isotype control antibody. The FCh distribution within each condition is represented using Waterfall plots. Changes between conditions in Waterfall plots were compared by linear mixed-effects model (formula: *Y∼condition_j_ + (1|sample))* using individuals as a random effect and the αPD-1 as an intra-group reference or Etvac as an inter-group reference. For cytokines, the multivariate distribution of samples and cytokines is represented as a Heatmap based on Euclidean distances.

## Acknowledgements

We thank the Flow Cytometry Core Facility at the German Trias I Pujol Research Institute (IGTP) and JC for comments on the initial draft and CM for study samples. This research was partially supported by the National Health Institute Carlos III grant PI17/00164. OBL was funded by an AGAUR-FI_B 00582 PhD fellowship from the Catalan Government and the European Social Fund. This study has received partial funding from Merck Sharp & Dohme LLC, a subsidiary of Merck & Co., Inc., Rahway, NJ, USA. The funders had no role in study design, data collection, analysis, or the decision to publish the manuscript.

## Author contributions

MM, AR, EJM and JGP conceptualized and designed the experiments. AR, EJM, MM, RP, OBL, DG and BH provided materials and performed and analysed some experiments. MM, DO, and JGP performed and revised statistical analyses. BM, CB, TH, BC, and JGP select study participants and samples. MM and JGP wrote the manuscript. All authors revised it critically for important intellectual content and have approved the final version submitted for publication.

## References

1. Deeks SG, Lewin SR, Havlir D V. The end of AIDS: HIV infection as a chronic disease. The Lancet. Published online 2013.

2. Rosenberg ES, Altfeld M, Poon SH, et al. Immune control of HIV-1 after early treatment of acute infection. Nature. Published online 2000. Doi:10.1038/35035103

3. Fischer M, Hafner R, Schneider C, et al. HIV RNA in plasma rebounds within days during structured treatment interruptions. AIDS. Published online 2003. Doi:10.1097/00002030-200301240-00009

4. Ta TM, Malik S, Anderson EM, et al. Insights Into Persistent HIV-1 Infection and Functional Cure: Novel Capabilities and Strategies. Front Microbiol. 2022;13:862270. Doi:10.3389/FMICB.2022.862270

5. Lichterfeld M, Gao C, Yu XG. An ordeal that does not heal: understanding barriers to a cure for HIV-1 infection. Trends Immunol. 2022;43(8):608–616. Doi:10.1016/j.it.2022.06.002

6. Perreau M, Banga R, Pantaleo G. Targeted Immune Interventions for an HIV-1 Cure. Trends Mol Med. Published online 2017. Doi:10.1016/j.molmed.2017.08.006

7. Cillo AR, Mellors JW. Which therapeutic strategy will achieve a cure for HIV-1? Curr Opin Virol. 2016;18:14–19. Doi:10.1016/j.coviro.2016.02.001

8. Murakoshi H, Akahoshi T, Koyanagi M, et al. Clinical Control of HIV-1 by Cytotoxic T Cells Specific for Multiple Conserved Epitopes. J Virol. Published online 2015. Doi:10.1128/jvi.00020-15

9. Hanke T. Aiming for protective T-cell responses: a focus on the first generation conserved-region HIVconsv vaccines in preventive and therapeutic clinical trials. Expert Rev Vaccines. 2019;18(10):1029–1041. Doi:10.1080/14760584.2019.1675518

10. Gilbert SC. T-cell-inducing vaccines – what’s the future. Immunology. Published online 2012. Doi:10.1111/j.1365-2567.2011.03517.x

11. Harty JT, Badovinac VP. Shaping and reshaping CD8+ T-cell memory. Nat Rev Immunol. Published online 2008. Doi:10.1038/nri2251

12. Collins DR, Gaiha GD, Walker BD. CD8+ T cells in HIV control, cure and prevention. Nature Reviews Immunology (2020). Doi:10.1038/s41577-020-0274-9

13. Colby DJ, Sarnecki M, Barouch DH, et al. Safety and immunogenicity of Ad26 and MVA vaccines in acutely treated HIV and effect on viral rebound after antiretroviral therapy interruption. Nature Medicine 2020 26:4. 2020;26(4):498–501. Doi:10.1038/S41591-020-0774-Y

14. Rosás-Umbert M, Mothe B, Noguera-Julian M, et al. Virological and immunological outcome of treatment interruption in HIV-1-infected subjects vaccinated with MVA-B. PloS One. 2017;12(9):e0184929. Doi:10.1371/JOURNAL.PONE.0184929

15. Mylvaganam GH, Silvestri G, Amara RR. HIV therapeutic vaccines: moving towards a functional cure. Curr Opin Immunol. 2015;35:1–8. Doi:10.1016/j.coi.2015.05.001

16. Schooley RT, Spritzler J, Wang H, et al. AIDS Clinical Trials Group 5197: A Placebo-Controlled Trial of Immunization of HIV-1-Infected Persons with a Replication-Deficient Adenovirus Type 5 Vaccine Expressing the HIV-1 Core Protein. J Infect Dis. 2010;202(5):705–716. Doi:10.1086/655468

17. Deeks SG, Archin N, Cannon P, et al. Research priorities for an HIV cure: International AIDS Society Global Scientific Strategy 2021. Nature Medicine 2021 27:12. 2021;27(12):2085–2098. Doi:10.1038/s41591-021-01590-5

18. Fuertes Marraco SA, Neubert NJ, Verdeil G, Speiser DE. Inhibitory Receptors Beyond T Cell Exhaustion. Front Immunol. 2015;6. Doi:10.3389/fimmu.2015.00310

19. Trautmann L, Janbazian L, Chomont N, et al. Upregulation of PD-1 expression on HIV-specific CD8+ T cells leads to reversible immune dysfunction. Nat Med. Published online 2006. Doi:10.1038/nm1482

20. Petrovas C, Casazza JP, Brenchley JM, et al. PD-1 is a regulator of virus-specific CD8+ T cell survival in HIV infection. Journal of Experimental Medicine. Published online 2006. Doi:10.1084/jem.20061496

21. Jones RB, Ndhlovu LC, Barbour JD, et al. Tim-3 expression defines a novel population of dysfunctional T cells with highly elevated frequencies in progressive HIV-1 infection. Journal of Experimental Medicine. Published online 2008. Doi:10.1084/jem.20081398

22. Hokey DA, Johnson FB, Smith J, et al. Activation drives PD-1 expression during vaccine-specific proliferation and following lentiviral infection in macaques. Eur J Immunol. Published online 2008. Doi:10.1002/eji.200737857

23. Allie SR, Zhang W, Fuse S, Usherwood EJ. Programmed death 1 regulates development of central memory CD8 T cells after acute viral infection. J Immunol. 2011;186(11):6280–6286. Doi:10.4049/JIMMUNOL.1003870

24. Sharpe AH, Pauken KE. The diverse functions of the PD1 inhibitory pathway. Nature Reviews Immunology 2017 18:3. 2017;18(3):153–167. Doi:10.1038/NRI.2017.108

25. Sehrawat S, Reddy PBJ, Rajasagi N, Suryawanshi A, Hirashima M, Rouse BT. Galectin-9/TIM-3 interaction regulates virus-specific primary and memory CD8 T cell response. PloS Pathog. 2010;6(5):1–16. Doi:10.1371/JOURNAL.PPAT.1000882

26. Day CL, Kaufmann DE, Kiepiela P, et al. PD-1 expression on HIV-specific T cells is associated with T-cell exhaustion and disease progression. Nature. 2006;443(7109):350–354. Doi:10.1038/nature05115

27. Chew GM, Fujita T, Webb GM, et al. TIGIT Marks Exhausted T Cells, Correlates with Disease Progression, and Serves as a Target for Immune Restoration in HIV and SIV Infection. PloS Pathog. 2016;12(1):e1005349. Doi:10.1371/JOURNAL.PPAT.1005349

28. Fromentin R, DaFonseca S, Costiniuk CT, et al. PD-1 blockade potentiates HIV latency reversal ex vivo in CD4+ T cells from ART-suppressed individuals. Nature Communications 2019 10:1. 2019;10(1):1–7. Doi:10.1038/s41467-019-08798-7

29. He X, Wong YC, Zhong M, et al. A follow-up study: 6-year cART-free virologic control of rhesus macaques after PD-1-based DNA vaccination against pathogenic SHIV SF162P3CN challenge. Microbiol Spectr. 2023;11(6). Doi:10.1128/SPECTRUM.03350-23

30. Finnefrock AC, Tang A, Li F, et al. PD-1 Blockade in Rhesus Macaques: Impact on Chronic Infection and Prophylactic Vaccination. The Journal of Immunology. Published online 2009. Doi:10.4049/jimmunol.182.2.980

31. Rahman SA, Yagnik B, Bally AP, et al. PD-1 blockade and vaccination provide therapeutic benefit against SIV by inducing broad and functional CD8+ T cells in lymphoid tissue. Sci Immunol. 2021;6(63). Doi:10.1126/SCIIMMUNOL.ABH3034

32. Bagchi S, Yuan R, Engleman EG. Immune Checkpoint Inhibitors for the Treatment of Cancer: Clinical Impact and Mechanisms of Response and Resistance. Annu Rev Pathol. 2021;16:223–249. Doi:10.1146/ANNUREV-PATHOL-042020-042741

33. Waldman AD, Fritz JM, Lenardo MJ. A guide to cancer immunotherapy: from T cell basic science to clinical practice. Nature Reviews Immunology 2020 20:11. 2020;20(11):651-668. Doi:10.1038/s41577-020-0306-5

34. Mutua G, Farah B, Langat R, et al. Broad HIV-1 inhibition in vitro by vaccine-elicited CD8+ T cells in African adults. Mol Ther Methods Clin Dev. 2016;3:16061. Doi:10.1038/mtm.2016.61

35. Mothe B, Manzardo C, Sanchez-Bernabeu A, et al. Therapeutic Vaccination Refocuses T-cell Responses Towards Conserved Regions of HIV-1 in Early Treated Individuals (BCN 01 study). EclinicalMedicine. 2019;11(June):65–80. Doi:10.1016/j.eclinm.2019.05.009

36. Szklarczyk D, Gable AL, Lyon D, et al. STRING v11: Protein-protein association networks with increased coverage, supporting functional discovery in genome-wide experimental datasets. Nucleic Acids Res. 2019;47(D1):D607–D613. Doi:10.1093/nar/gky1131

37. Velu V, Titanji K, Zhu B, et al. Enhancing SIV-specific immunity in vivo by PD-1 blockade. Nature. 2009;458(7235):206–210. Doi:10.1038/nature07662

38. Harper J, Gordon S, Chan CN, et al. CTLA-4 and PD-1 dual blockade induces SIV reactivation without control of rebound after antiretroviral therapy interruption. Nat Med. 2020;26(4):519–528. Doi:10.1038/S41591-020-0782-Y

39. Mothe B, Rosás-Umbert M, Coll P, et al. HIVconsv Vaccines and Romidepsin in Early-Treated HIV-1-Infected Individuals: Safety, Immunogenicity and Effect on the Viral Reservoir (Study BCN02). Front Immunol. 2020;11:823. Doi:10.3389/FIMMU.2020.00823

40. Uldrick TS, Adams S V., Fromentin R, et al. Pembrolizumab induces HIV latency reversal in people living with HIV and cancer on antiretroviral therapy. Sci Transl Med. 2022;14(629):3836. Doi:10.1126/scitranslmed.abl3836

41. Moyo N, Borthwick NJ, Wee EG, et al. Long-term follow up of human T-cell responses to conserved HIV-1 regions elicited by DNA/simian adenovirus/MVA vaccine regimens. PloS One. 2017;12(7):1–19. Doi:10.1371/journal.pone.0181382

42. Harper J, Ribeiro SP, Chan CN, et al. Interleukin-10 contributes to reservoir establishment and persistence in SIV-infected macaques treated with antiretroviral therapy. J Clin Invest. 2022;132(8). Doi:10.1172/JCI155251

43. Gonzalez-Cao M, Morán T, Dalmau J, et al. Assessment of the Feasibility and Safety of Durvalumab for Treatment of Solid Tumors in Patients With HIV-1 Infection: The Phase 2 DURVAST Study. JAMA Oncol. 2020;6(7):1063–1067. Doi:10.1001/JAMAONCOL.2020.0465

44. Blanch-Lombarte O, Gálvez C, Revollo B, et al. Enhancement of Antiviral CD8+ T-Cell Responses and Complete Remission of Metastatic Melanoma in an HIV-1-Infected Subject Treated with Pembrolizumab. J Clin Med. 2019;8(12). Doi:10.3390/JCM8122089

45. Ribas A, Wolchok JD. Cancer immunotherapy using checkpoint blockade. Science (1979). Published online 2018. Doi:10.1126/science.aar4060

46. Gonzalez-Cao M, Martinez-Picado J, Karachaliou N, Rosell R, Meyerhans A. Cancer immunotherapy of patients with HIV infection. Clinical and Translational Oncology. 2019;21(6):713–720. Doi:10.1007/S12094-018-1981-6

47. Uldrick TS, Ison G, Rudek MA, et al. Modernizing clinical trial eligibility criteria: Recommendations of the American society of clinical oncology-friends of cancer research HIV working group. Journal of Clinical Oncology. 2017;35(33):3774–3780. Doi:10.1200/JCO.2017.73.7338

48. Xu-Monette ZY, Zhang M, Li J, Young KH. PD-1/PD-L1 blockade: Have we found the key to unleash the antitumor immune response? Front Immunol. 2017;8(DEC):1597. Doi:10.3389/FIMMU.2017.01597

49. Sun W, Gao C, Hartana CA, et al. Phenotypic signatures of immune selection in HIV-1 reservoir cells. Nature 2023 614:7947. 2023;614(7947):309–317. Doi:10.1038/s41586-022-05538-8

50. Banga R, Procopio FA, Noto A, et al. PD-1(+) and follicular helper T cells are responsible for persistent HIV-1 transcription in treated aviremic individuals. Nat Med. 2016;22(7):754–761. Doi:10.1038/NM.4113

51. Bailón L, Llano A, Cedeño S, et al. Safety, immunogenicity and effect on viral rebound of HTI vaccines in early treated HIV-1 infection: a randomized, placebo-controlled phase 1 trial. Nature Medicine 2022 28:12. 2022;28(12):2611–2621. Doi:10.1038/s41591-022-02060-2

52. Beatriz Mothe Pujades, Adrià Curran, Juan Carlos López, et al. A PLACEBO-CONTROLLED RANDOMIZED TRIAL OF THE HTI IMMUNOGEN VACCINE AND VESATOLIMOD - Abstract Number 433. In: CROI Conference. ; 2023.

53. Michael J.Peluso, Amelia Deitchman, Gesham Magombedze, et al. REBOUND DYNAMICS FOLLOWING IMMUNOTHERAPY WITH AN HIV VACCINE, TLR9 AGONIST, AND bNAbs - Abstract Number 435. In: CROI Conference. ; 2023.

54. Julg B, Stephenson KE, Tomaka F, et al. Immunogenicity of 2 therapeutic mosaic HIV-1 vaccine strategies in individuals with HIV-1 on antiretroviral therapy. Npj Vaccines 2024 9:1. 2024;9(1):1–10. Doi:10.1038/s41541-024-00876-2

55. Létourneau S, Im EJ, Mashishi T, et al. Design and Pre-Clinical Evaluation of a Universal HIV-1 Vaccine. PloS One. 2007;2(10):e984. Doi:10.1371/JOURNAL.PONE.0000984

56. Wu G, Swanson M, Talla A, et al. HDAC inhibition induces HIV-1 protein and enables immune-based clearance following latency reversal. JCI Insight. Published online 2017. Doi:10.1172/jci.insight.92901

57. Passaes CPB, Bruel T, Decalf J, et al. Ultrasensitive HIV-1 p24 Assay Detects Single Infected Cells and Differences in Reservoir Induction by Latency Reversal Agents. J Virol. Published online 2017. Doi:10.1128/jvi.02296-16

58. Van Der Maaten L, Hinton G. Visualizing Data using t-SNE. Journal of Machine Learning Research. 2008;9:2579–2605.

59. Cho H, Berger B, Peng J. Generalizable and Scalable Visualization of Single-Cell Data Using Neural Networks. Cell Syst. 2018;7(2):185–191.e4. doi:10.1016/J.CELS.2018.05.017

60. Levine JH, Simonds EF, Bendall SC, et al. Data-Driven Phenotypic Dissection of AML Reveals Progenitor-like Cells that Correlate with Prognosis. Cell. Published online 2015. Doi:10.1016/j.cell.2015.05.047

61. Blondel VD, Guillaume JL, Lambiotte R, Lefebvre E. Fast unfolding of communities in large networks. Journal of Statistical Mechanics: Theory and Experiment. Published online 2008. Doi:10.1088/1742-5468/2008/10/P10008

62. R Development Core Team. R: A language and environment for statistical computing. *Vienna, Austria*. Published online 2017. Doi:R Foundation for Statistical Computing, Vienna, Austria. ISBN 3-900051-07-0, URL http://www.R-project.org.

