## Supplemental Figures for "PD-1 blockade enhances HIV-1 vaccine-induced CD8⁺ T-cell responses in PWH early ART-treated"

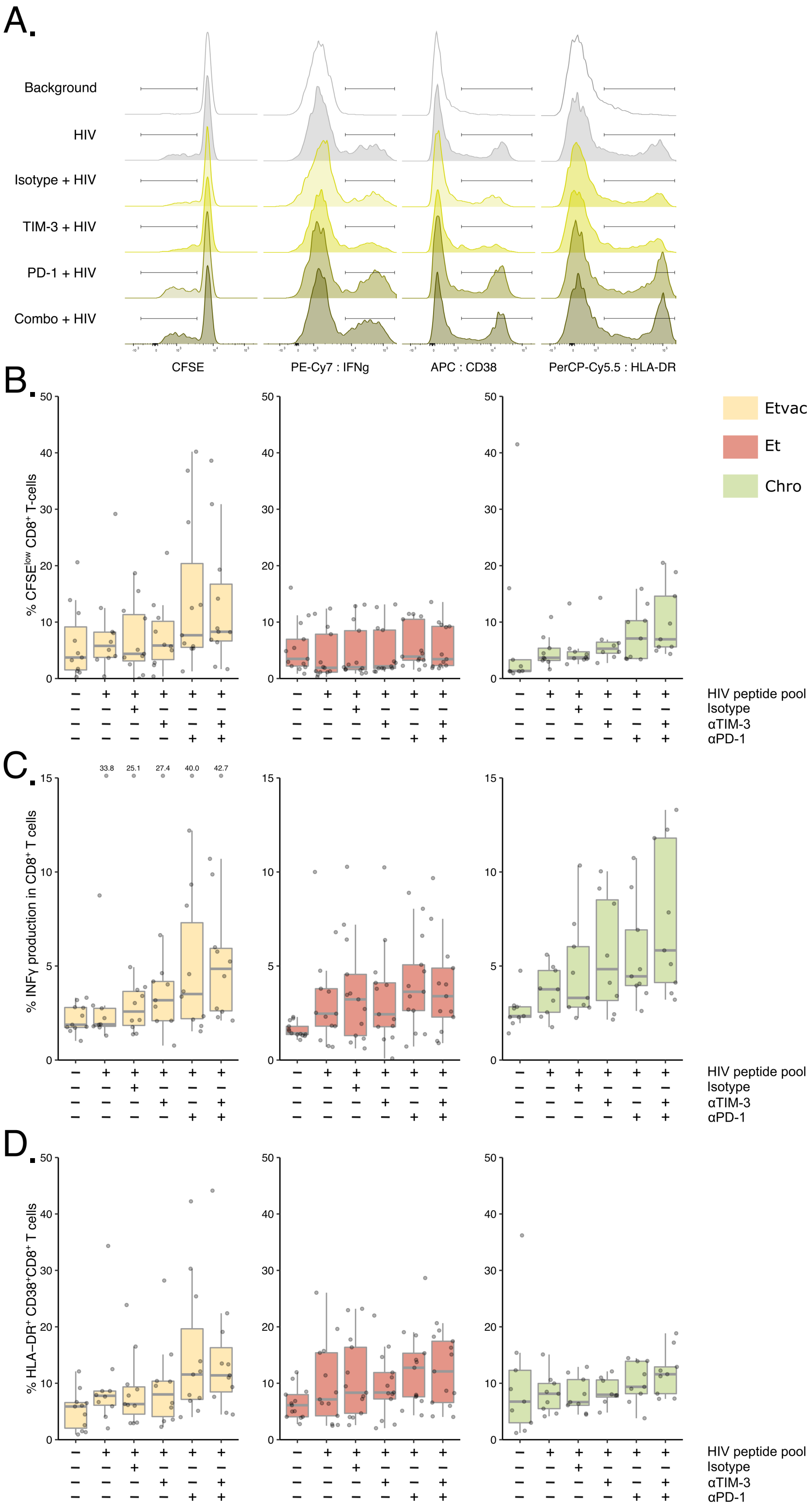


**Figure Supplementary 1. Proliferation, IFNγ and HLA-DR$^+$CD38$^+$ expression in total CD8$^+$ T cells.** **A** Representative histogram from an Etvac individual (Etvac11). Overlaid histograms show the modal distribution of CFSE, IFNγ production and expression of CD38 and HLA-DR in the CD8$^+$ T cell population in the different experimental conditions. **B** Boxplot showing proliferation (CFSE$^{low}$), IFNγ production and co-expression of HLA-DR$^+$CD38$^+$ in total CD8$^+$ T cells in the three study groups. To estimate the HIV-1-specific fraction, we subtracted the CFSE loss, IFNγ production and co-expression of HLA-DR$^+$CD38$^+$ in the absence of the corresponding HIV peptide pool.

A.

**BIOLOGICAL PROCESS**

| | GO-term | Description | Count in gene set | false discovery rate |
|---|---|---|---|---|
| 1 | GO:0006955 | immune response | 13 of 1560 | 6.80e-12 |
| 2 | GO:0019221 | cytokine-mediated signaling pathway | 10 of 655 | 3.06e-10 |
| 3 | GO:0010469 | regulation of signaling receptor activity | 9 of 577 | 3.57e-09 |
| 4 | GO:0051707 | response to other organism | 9 of 835 | 5.28e-08 |
| 5 | GO:0046427 | positive regulation of receptor signaling pathway via JAK-STAT | 5 of 73 | 1.36e-07 |
| 6 | GO:0042981 | regulation of apoptotic process | 10 of 1501 | 1.72e-07 |
| 7 | GO:0070201 | regulation of establishment of protein localization | 8 of 662 | 1.75e-07 |
| 8 | GO:1904951 | positive regulation of establishment of protein localization | 7 of 397 | 1.91e-07 |
| 9 | GO:0050708 | regulation of protein secretion | 7 of 422 | 2.71e-07 |
| 10 | GO:0006952 | defense response | 9 of 1234 | 5.96e-07 |

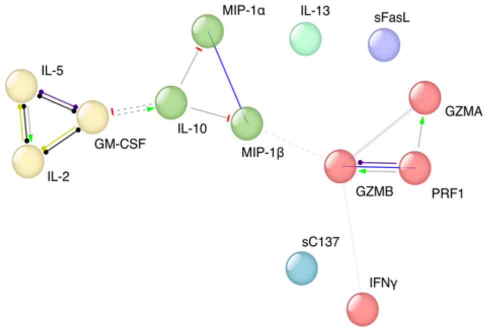


B.

**KEGG PATHWAYS**

| | Pathway | Description | Count in gene set | false discovery rate |
|---|---|---|---|---|
| 1 | hsa04060 | Cytokine-cytokine receptor interaction | 10 of 263 | 4.03e-15 |
| 2 | hsa05330 | Allograft rejection | 7 of 35 | 6.50e-15 |
| 3 | hsa05320 | Autoimmune thyroid disease | 6 of 49 | 1.30e-11 |
| 4 | hsa05332 | Graft-versus-host disease | 5 of 36 | 6.16e-10 |
| 5 | hsa04940 | Type I diabetes mellitus | 5 of 40 | 8.03e-10 |
| 6 | hsa05321 | Inflammatory bowel disease (IBD) | 5 of 62 | 5.25e-09 |
| 7 | hsa04630 | Jak-STAT signaling pathway | 6 of 160 | 5.25e-09 |
| 8 | hsa05142 | Chagas disease (American trypanosomiasis) | 5 of 101 | 3.70e-08 |
| 9 | hsa04660 | T cell receptor signaling pathway | 5 of 99 | 3.70e-08 |
| 10 | hsa04650 | Natural killer cell mediated cytotoxicity | 5 of 124 | 8.78e-08 |

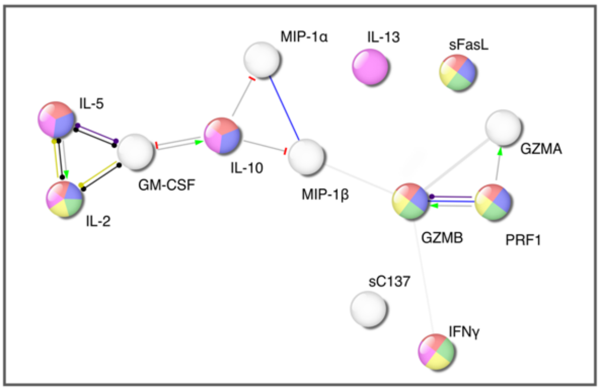


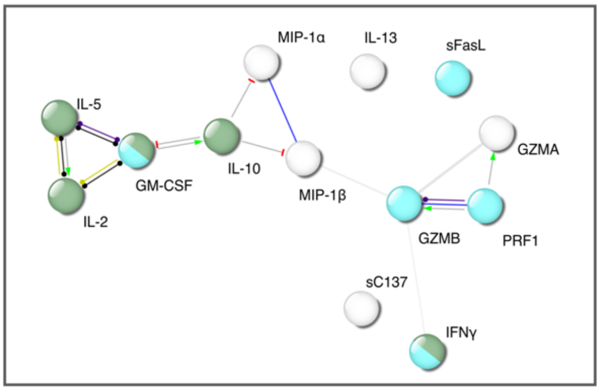


**Figure Supplementary 2. Enrichment functional analysis by STRING. A.** Cytokine network and list of biological processes related with the cytokines detected in response to αPD-1 **B**. KEGG pathways in which PD-1-detected cytokines are implicated.
